# The impact of antigenic escape on the evolution of virulence

**DOI:** 10.1101/2021.01.19.427227

**Authors:** Akira Sasaki, Sébastien Lion, Mike Boots

## Abstract

Understanding the evolutionary drivers determining the transmission rate and virulence of pathogens remains an important challenge for evolutionary theory with clear implications to the control of human, agricultural and wildlife infectious disease. Although disease is often very dynamic, classical theory examines the long-term outcome of evolution at equilibrium and, in simple models, typically predicts that R_0_ is maximized. For example, immune escape may lead to complex disease dynamics including repeated epidemics, fluctuating selection and diversification. Here we model the impact of antigenic drift and escape on the evolution of virulence and show analytically that these non-equilibrium dynamics select for more acute pathogens with higher virulence. Specifically, under antigenic drift and when partial cross immunity leads to antigenic escape, our analysis predicts the long-term maximization of the intrinsic growth rate of the parasite resulting in more acute and virulent pathogens than those predicted by classic R_0_ maximization. Furthermore, it follows that these pathogens will have a lower R_0_ leading to implications for epidemic, endemic behavior and control. Our analysis predicts both the timings and outcomes of antigenic shifts leading to repeated epidemics and predicts the increase in variation in both antigenicity and virulence before antigenic escape. There is considerable variation in the degree of antigenic escape that occurs across pathogens and our results may help to explain the difference in virulence between related pathogens most clearly seen in the human A, B and C influenzas. More generally our results show the importance of examining the evolutionary consequences of non-equilibrium dynamics.

## Introduction

Recent epidemics emphasize that infectious disease remains a major problem for human health and agriculture [1–4] and they are increasingly recognized as important in ecosystems and conservation [5,6]. This importance has led to the development of extensive theoretical literature on the epidemiology, ecology and evolution of host-pathogen interactions [7–10]. Understanding the drivers of the evolution of virulence, typically defined in the evolutionary literature as the increased death rate of individuals due to infection, is a key motivator of this theoretical work [8,10–14]. Generally, models assume that a higher transmission rate trade-offs against the intrinsic cost of reducing the infectious period due to higher death rates (virulence) of rapidly dividing strains and classically predict that evolution maximizes the parasite epidemiological R_0_ leading to an optimal virulence [8,10–14]. In fact, this result only holds in models where ecological feedbacks take a constrained form, such that even relatively simple processes such as density-dependent mortality, multiple infections and spatial structure may lead to diversification or different optima [10,12,13,15]. Moreover, this classic evolutionary theory examines the long-term equilibrium evolutionary outcome in the context of stable endemic diseases, but in nature infectious diseases often exhibit complex dynamics, with potentially important impacts on pathogen fitness [16–19].

Antibody-mediated immunity is a critical factor driving the dynamics of important infectious diseases such as seasonal influenza, leading to selection for novel variants that can escape immunity to the current predominant strain [20–22]. Such antigenic escape typically causes the optimal strain of the parasite to change through time as it moves through antigenic space. Moreover, partial cross-immunity between the different parasite strains may lead to recurrent epidemics, fluctuations in parasite strains and potentially strain coexistence [23–26]. Current theory has shown that the evolution of immune escape can lead to dramatic disease outbreaks [24–26], but the implications of these epidemiological dynamics for the evolution of disease virulence are unknown. This question is challenging in part because much of the theoretical framework used to study virulence evolution typically considers diseases that are at an endemic equilibrium [8,10–14]. As such we currently lack a broad theoretical understanding of the evolution of virulence in the presence of antigenic escape, despite its importance as an epidemic process and the likely implications of its inherently dynamical epidemic nature.

Here we examine the implications of antigenic escape for the evolution of infectious disease in the context of the well-studied transmission/virulence trade-off [10,27]. For simplicity we first examine analytically the case without cross-immunity and then apply a new ‘oligomorphic’ analysis that combines quantitative genetic and game theoretical approaches [28] to examine the impact of cross-immunity that leads to antigenic jumps and epidemic outbreaks. With this analysis we are able to study the evolutionary outcome across a range of ecological and evolutionary time scales allowing us to examine evolutionary outcomes under non-equilibrium conditions. Our key result is that antigenic escape selects for higher transmission and virulence due to the repeated epidemics caused by immune escape, leading to the long-term persistence of acute pathogens. Indeed, antigenic escape has the potential to select for infectious diseases with substantially higher virulence than that predicted by the maximization of R_0_ in classic disease models leading to the evolution of much more acute diseases.

## Modeling

We consider a population of pathogens structured by a one-dimensional antigenic trait *x*, so that *I*(*t, x*) is the density of hosts infected with antigenicity strain *x* at time *t*. Following Gog and Grenfell [29], we assume that an individual is either perfectly susceptible or perfectly immune to a strain. A strain of pathogen can infect any host, but will be infectious only when the host is susceptible to that strain. When a strain *y* of pathogen infects a host that is susceptible to a strain *x*, the host may become (perfectly) immune to the strain *x* with probability *σ*(*x* – *y*). This is the partial cross immunity function between strain *x* and *y*, that takes a value between 0 and 1 and is a decreasing function of antigenic distance |*x* – *y*| between strain *x* and *y*. The density of hosts susceptible to antigenicity strain *x* at time *t* is noted *S*(*t, x*).

Assuming that all pathogen strains have the same transmission rate *β* and virulence *α*, we can describe the dynamics with the following structured Susceptible-Infected-Recovered model:

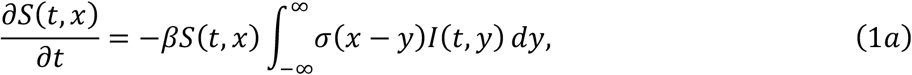

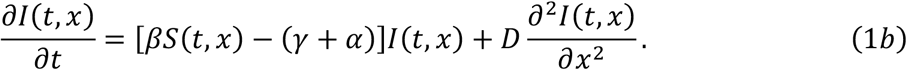

where *γ* is the recovery rate, and 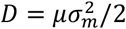 is one half of the mutation variance for the change in antigenicity, representing random mutation in the continuous antigenic space. The dynamics for the density of recovered hosts is omitted from (1) as it does not affect the dynamics (1) for the densities of susceptible and infected hosts.

### Invasion of a single pathogen

In our first scenario, we start with a population where all hosts are susceptible to any strain (*S*(0, *x*) = 1) and a pathogen strain with antigenicity trait *x* = 0 is initially introduced, so that *I*(0, *x*) = *ϵδ*(*x*), where ***ϵ*** is a small positive constant and *δ*(·) is Dirac’s delta function. The system then exhibits travelling wave dynamics in antigenicity space. At the front of the travelling wave, *I*(*t, x*) is sufficiently small and *S*(*t, x*) is sufficiently close to 1. Eq. (1a) can then be linearized as

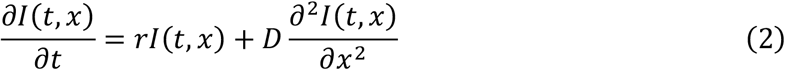

where *r* = *β* – (*γ* + *α*) is the rate of increase of an antigenicity strain before it spreads in the population and causes the build-up of herd immunity. The system (1) asymptotically approaches travelling waves of both pathogen antigenicity quasispecies *I*(*t, x*), which have an isolated peak around the current antigenicity, and host susceptibility profile *S*(*t, x*), that smoothly steps down towards a low level after pathogen antigenicity quasispecies passes through, with a common constant wave speed [30] (Fig 1A)

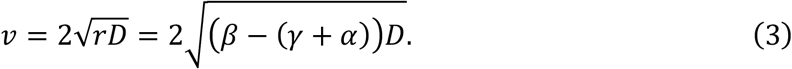

**Figure 1.**
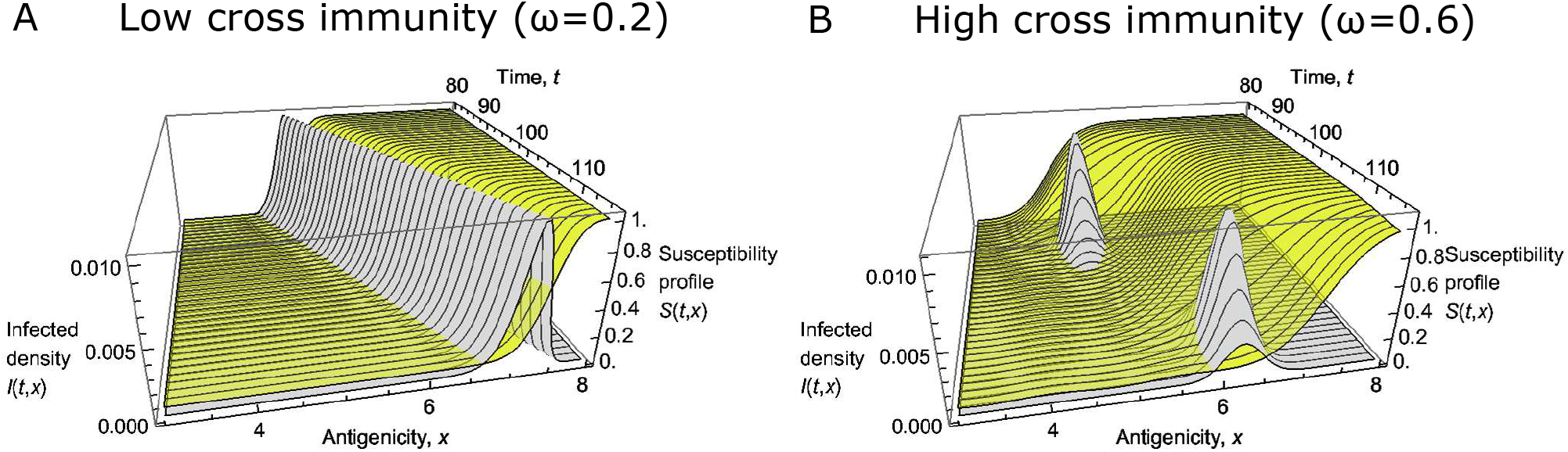
Continuous antigenic drift (A) and periodical antigenic shits (B) of the model. The orange colored surface denotes the infected density *I*(*t, x*) varying in time *t* and intigenicty *x*, and the yellow colored surface denotes the density of hosts *S*(*t, x*) that are susceptible to antigenicity strain *x* of pathogen at time *t*. The width of cross immunity is (*ω*) = 0.2 in (A) and *ω* = 0.6 in (B). Other parameters are *β* = 2, *u* = 0.001, *α* = 0.1, *γ* = 0.5, and *D* = 0.001.

As the width of the partial cross-immunity function *σ*(*x* – *y*) increases, the travelling wave with static shapes described above is destabilized (Fig #B, C), and the system shows intermittent outbreaks that occur periodically both in time and in antigenicity space [29,31] (Fig 1B). However, the wave speed is unchanged from (3), as the linearized system (2) towards the frontal end remains the same irrespective of the stability of wave profile that lags behind (Fig S1).

### Evolution of antigenic escape with cross-immunity

To predict how cross-immunity affects the evolution of antigenic escape, we use an oligomorphic dynamics analysis [28]. We consider a population composed of different antigenicity strains (or morphs), that can be viewed as quasispecies. The analysis in Appendix A allows us to track the dynamics of morph frequencies, *p_i_*, and mean trait values, 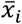, as:

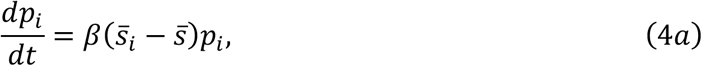

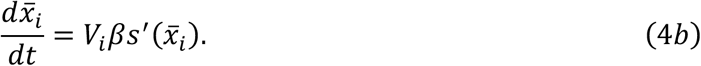

where *s*(*x*) is the susceptibility profile of the population, which depends on the cross-immunity function *σ*, 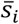 is the mean susceptibility perceived by viral morph *i*, and 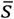 the mean susceptibility averaged over the different viral morphs. Note that, in general, *s*(*x*), 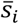 and 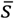 will be functions of time, as the susceptibility profile is molded by the epidemiological dynamics of *S*(*t, x*) and *I*(*t, x*).

Equation (4a) reveals that, as expected, morph *i* will increase in frequency if the susceptibility of the host population to this strain is higher on average. Equation (4b) shows that the increase in the mean antigenicity trait of morph *i* depends on (i) the variance of the morph distribution, *V_t_*, (ii) the transmission rate, and (iii) the slope of the susceptibility profile close to the morph mean 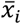. Together with an equation for the dynamics of variance under mutation and selection (SI: Appendix S1), equations (4a) and (4b) allow us to quantitatively predict the change in antigenicity after a primary outbreak, as shown in Figure 2.

**Figure 2.**
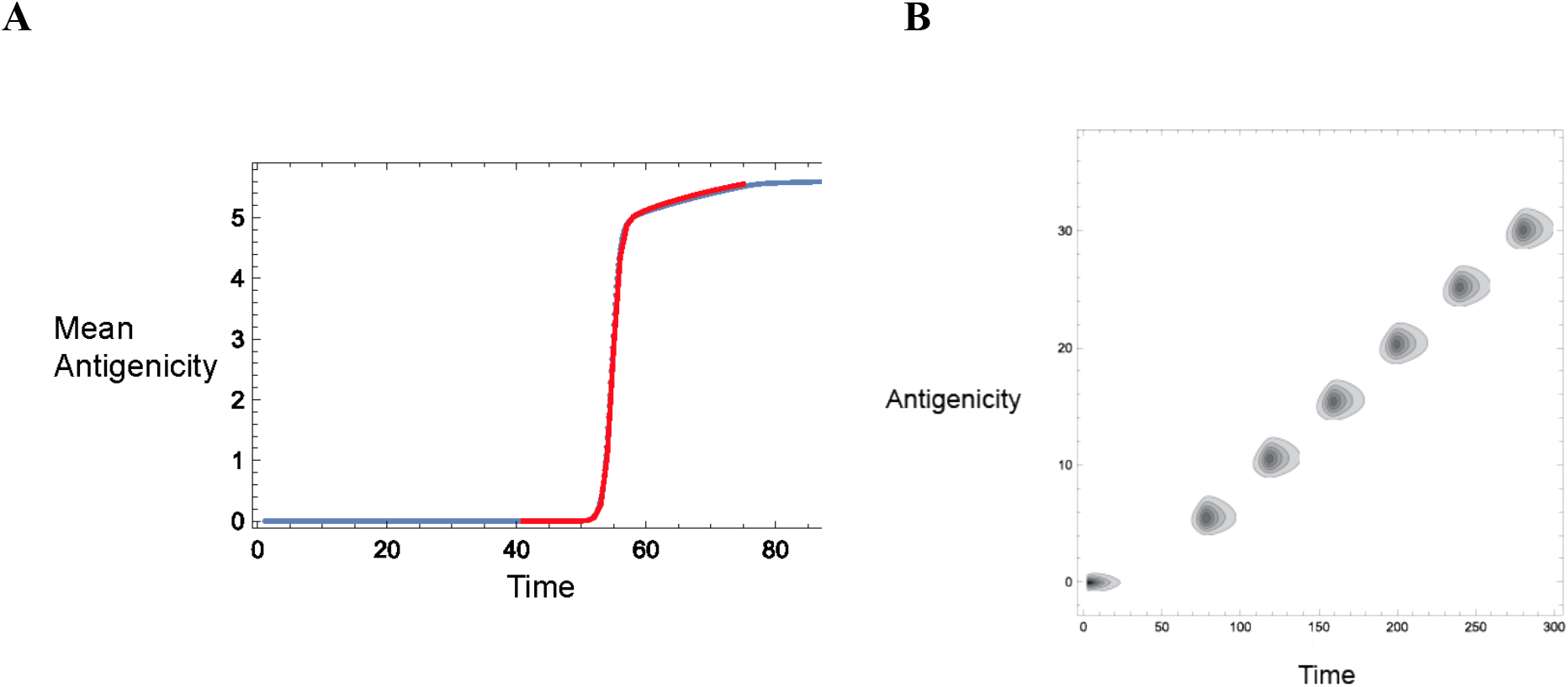
Oligomorphic dynamics prediction of the emergence of antigenicity shift. (a) Oligomorphic prediction for the change in the mean antigenicity after the primary outbreak at *x* = 0 (red curve), compared with that obtained by numerical simulations (blue dots). (b) Heat map representation of the time change of the antigenic drift model (1). Parameters: *β* = 2, *γ* + *α* = 0.6, *σ*(*x*) = exp(−*x*^2^/2*ω*^2^) with *ω* = 2, *D* = 0.001. Initially, all hosts are equally susceptible with *S*(0, *x*) = 1. The primary pathogen strain is introduced at *x* = 0 with infected density 0.001.

For instance, after a primary outbreak caused by a strain with antigenicity 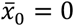 at *t* = 0, the susceptibility profile is approximately constant and given by *s*(*x*) = (1 – *ψ*_0_)^*σ*(*x*)^, where *ψ*_0_ is the final size of the epidemic of the primary outbreak at antigenicity *x* = 0 (SI: Appendix S1). Thus, for a decreasing cross-immunity function, *σ*(*x*), the slope of the susceptibility profile is positive, which selects for increased values of the mean antigenicity trait 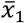, of a second emerging morph (SI: Appendix S1). As the process repeats itself, this leads to successive jumps in antigenic space. In addition, a more peaked cross-immunity function, *σ*, yields larger slopes to the susceptibility profile and thus selects for higher values of the antigenicity trait.

### Long-term joint evolution of antigenicity, transmission and virulence

We now extend our analysis to account for mutations affecting pathogen life-history traits such as transmission and virulence. To simplify, we use the classical assumption of a transmission-virulence trade-off [8,10–14] and consider that a pathogen morph, *i*, has frequency, *p_i_*, mean antigenicity trait, 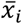, and mean virulence 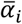. In Appendix S2, we show that the morph’s mean traits change as

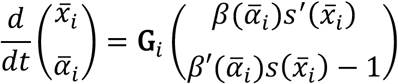

where **G**_*i*_ is the genetic (co)variance matrix, and the vector on the right-hand side is the selection gradient. Note that, while the selection gradient on antigenicity depends on the slope of the antigenicity profile at the morph mean, the selection gradient on virulence depends on the slope of the transmission-virulence trade-off at the morph mean, weighted by the susceptibility profile at the morph mean.

Assuming we can neglect the build-up of correlations between antigenicity and virulence due to mutation and selection, the genetic (co)variance matrix is diagonal with elements 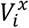 and 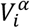. Then, as shown previously, antigenicity increases if the slope of the susceptibility profile is locally positive, while mean virulence increases as long as 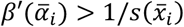. For a fixed antigenicity trait, *x* = *x*\*, the susceptibility profile converges towards *s*(*x*\*) = (*γ* + *α*)/*β* and the evolutionary endpoint satisfies

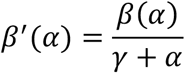

which corresponds to the classical result of *R*_0_ maximisation for the unstructured SI model [15,27]. However, when antigenicity can evolve, selection will also lead to the build-up of a positive covariance *C* between antigenicity and virulence, resulting in a synergistic effect (SI: Appendix S2; Figure S3). As the antigenicity trait increases, the evolutionary trajectory of virulence converges to the solution of

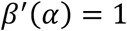

which corresponds to maximizing the rate of increase of pathogen *r*(*α*) = *β*(*α*) – (*γ* + *α*) in a fully susceptible population. This is equivalent to maximizing the wave speed 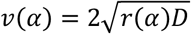, as shown in Appendix S3. Figure 3a shows that, in the absence of cross-immunity, the ES virulence is well predicted by r maximization. With cross-immunity (Figure 3b), virulence evolution is characterized by jumps that reflect the sudden shifts in antigenicity due to cross-immunity.

**Figure 3.**
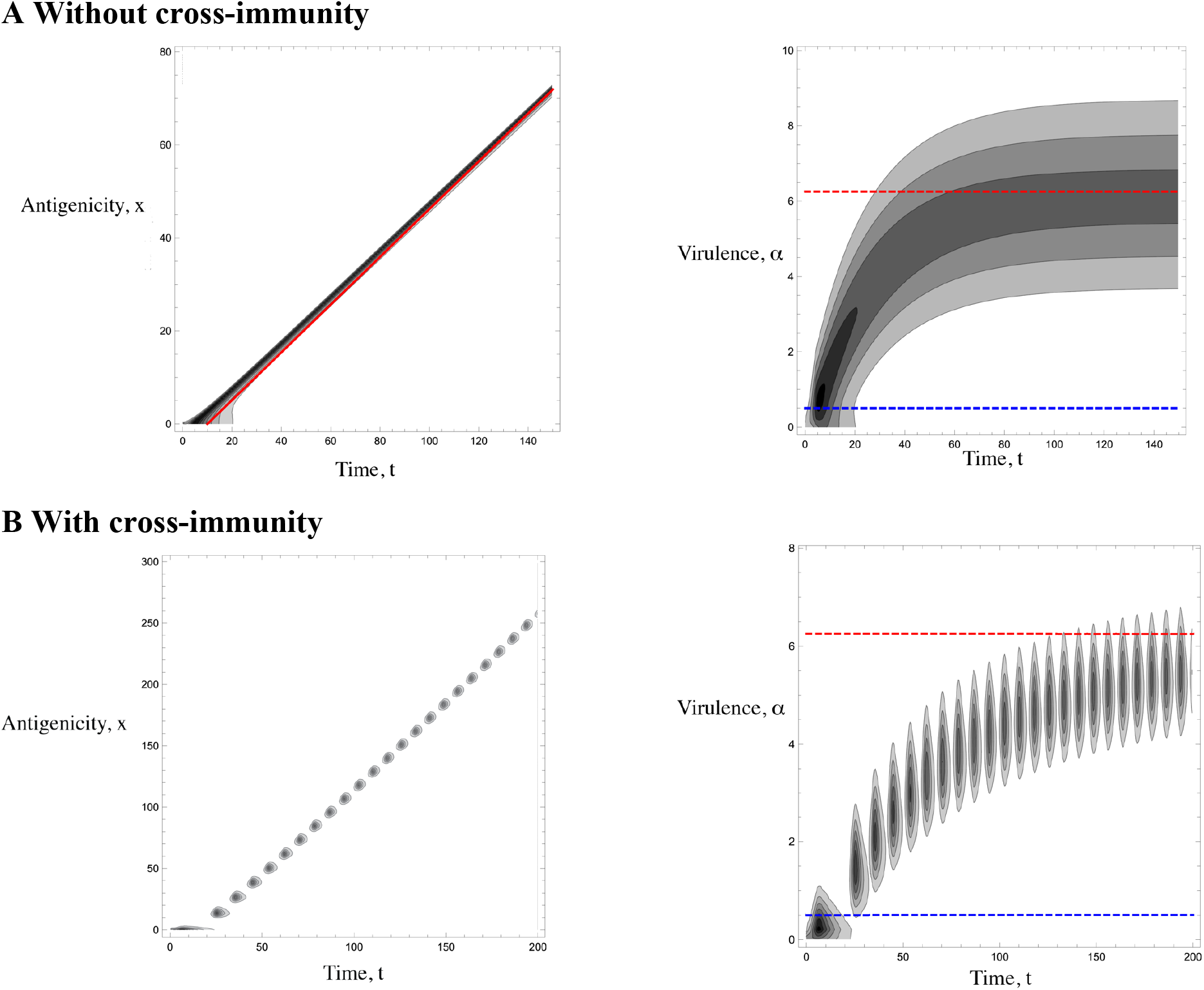
Marginal distribution of antigenicity (left) and virulence (right) in the absence (a) or presence (b) of cross-immunity. (a) No cross immunity is assumed so that each antigenicity genotype causes specific herd immunity: *σ*(*x* – *y*) = *δ*(*x* – *y*), where *δ*(·) is Dirac’s delta function. There are 1600 antigenic variations having equally divided antigenicity between 0 and 80. (b) We assume a Gaussian cross-immunity kernel, *σ*(*x* – *y*) = exp(−(*x* – *y*)^2^/2*ω*^2^), with width *ω* = 5. There are 300 antigenic variations having equally divided antigenicity between 0 and 300. In both panels, there are 100 viral virulence traits each having virulence equally divided between 0 and 20, and Diffusion constants due to mutations are *D_x_* = 0.01 (for antigenicity) and *D_α_* = 0.01(for virulence). The blue dashed lines show the ES virulence predicted from maximizing R_0_, as expected in the absence of antigenic escape, while the red dashed lines show the predicted ES virulence that maximizes *r*(*α*) = *β*(*α*) – (*γ* + *α*) as predicted from our analysis. Other parameters: *γ* = 0.5, and 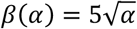.

As such antigenic escape selects for higher transmission and virulence and more acute infectious diseases. This has parallels with the results that show that there is a transient increase in virulence at the start of an epidemic with r rather than R_0_ being maximized [16,17,19,32], but it is important to note that here we predict the long-term persistence of highly transmissible and virulent disease strains due to the existence of antigenic escape.

Although we have so far assumed a never-ending antigenic escape process, it is easy to extend our analysis to consider that antigenic escape is constrained by pleiotropic effects. Then, once the antigenicity trait has stabilized, the ES virulence would satisfy

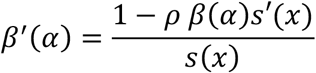

where *ρ* = *C/V_α_* measures the correlation between antigenicity and virulence. Thus, the slope to the transmission-virulence trade-off at the ESS now takes an intermediate value between *β*/(*γ* + *α*) and 1, as shown in figure 4.

**Figure 4.**
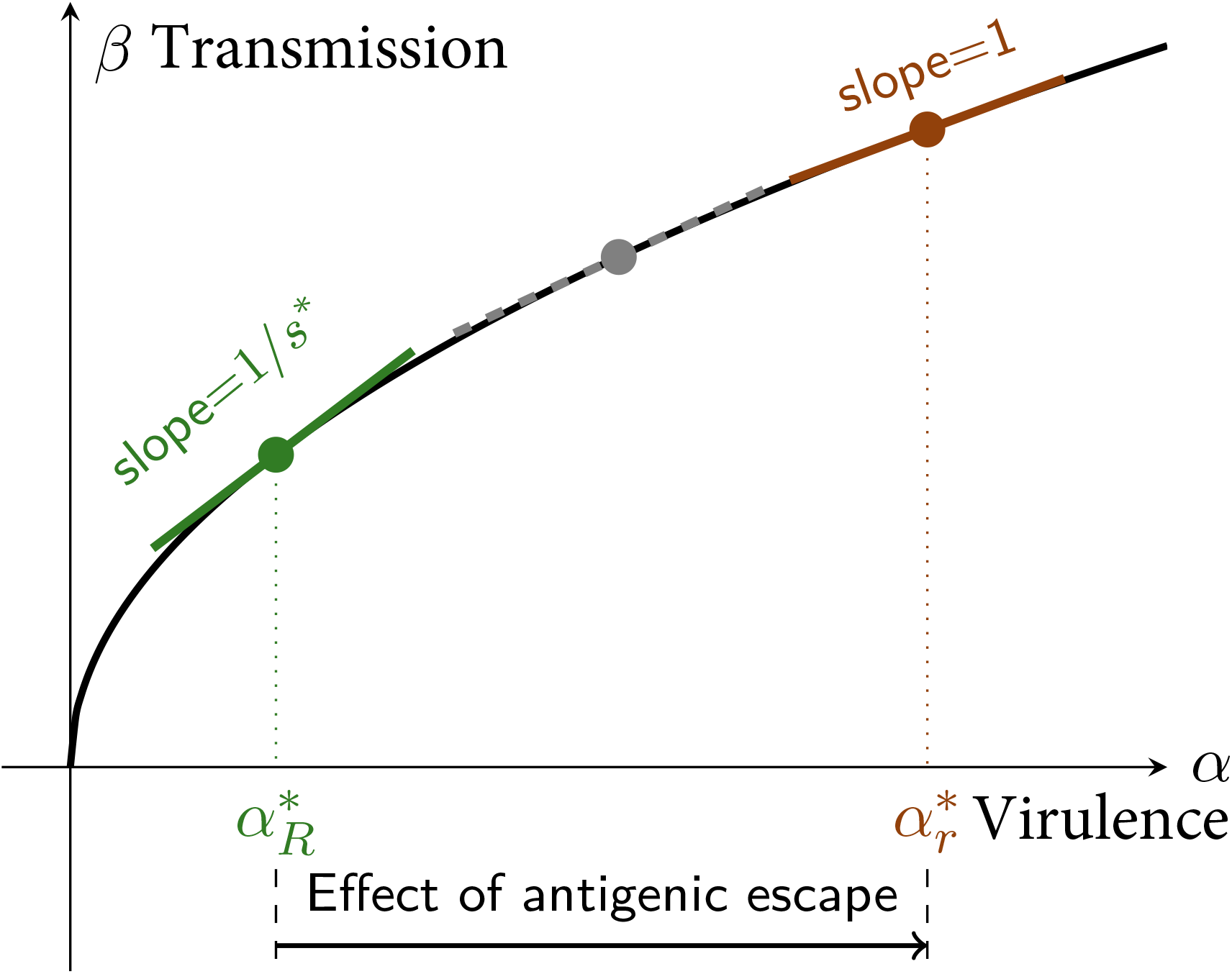
Graphical representation of the predicted ES virulence with or without antigenic escape under the assumption of a transmission-virulence trade-off. In the absence of antigenic escape, the ES virulence, 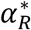, can be predicted from the maximization of the pathogen’s epidemiological basic reproduction ratio, *R*_0_ = *β*/(*γ* + *α*), or equivalently by the minimization of the total density of susceptible hosts since, at equilibrium, *s*\* = 1/*R*_0_. The slope of the transmission-virulence trade-off at the ESS is then 1/*s*\* = *R*_0_. With antigenic escape, the ES virulence, 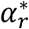, can be predicted from the maximization of the pathogen’s growth rate in a fully susceptible population, *r*_0_ = *β*(*α*) – (*γ* + *α*). The slope of the transmission-virulence trade-off at the ESS is then 1. This holds true in the limit of a large antigenicity trait, but intermediate values of ES virulence, corresponding to intermediate slopes, can also be selected for if other processes constrain the evolution of the antigenicity trait, as explained in the main text.

### Short-term joint evolution of antigenicity and virulence

Although our analysis allows us to understand the long-term evolution of pathogen traits, it can also be used to accurately predict the short-term dynamics of antigenicity and virulence. We now consider that a primary outbreak has molded a susceptibility profile *s*(*x*) that we assume constant. Although this assumption will cause deviations from the true susceptibility profile, it allows us to decouple our evolutionary oligomorphic dynamics from the epidemiological dynamics. Figure 5 shows that the approximation accurately predicts the jump in antigenic space and joint increase in virulence during the secondary outbreak. The accuracy of the prediction depends on the time at which we seed the oligomorphic dynamical system, as detailed in Appendix S1, but remains high for a broad range of values of this initial time. Hence, our analysis can be used to successfully predict the trait dynamics after the emergence of a new antigenic strain.

**Figure 5.**
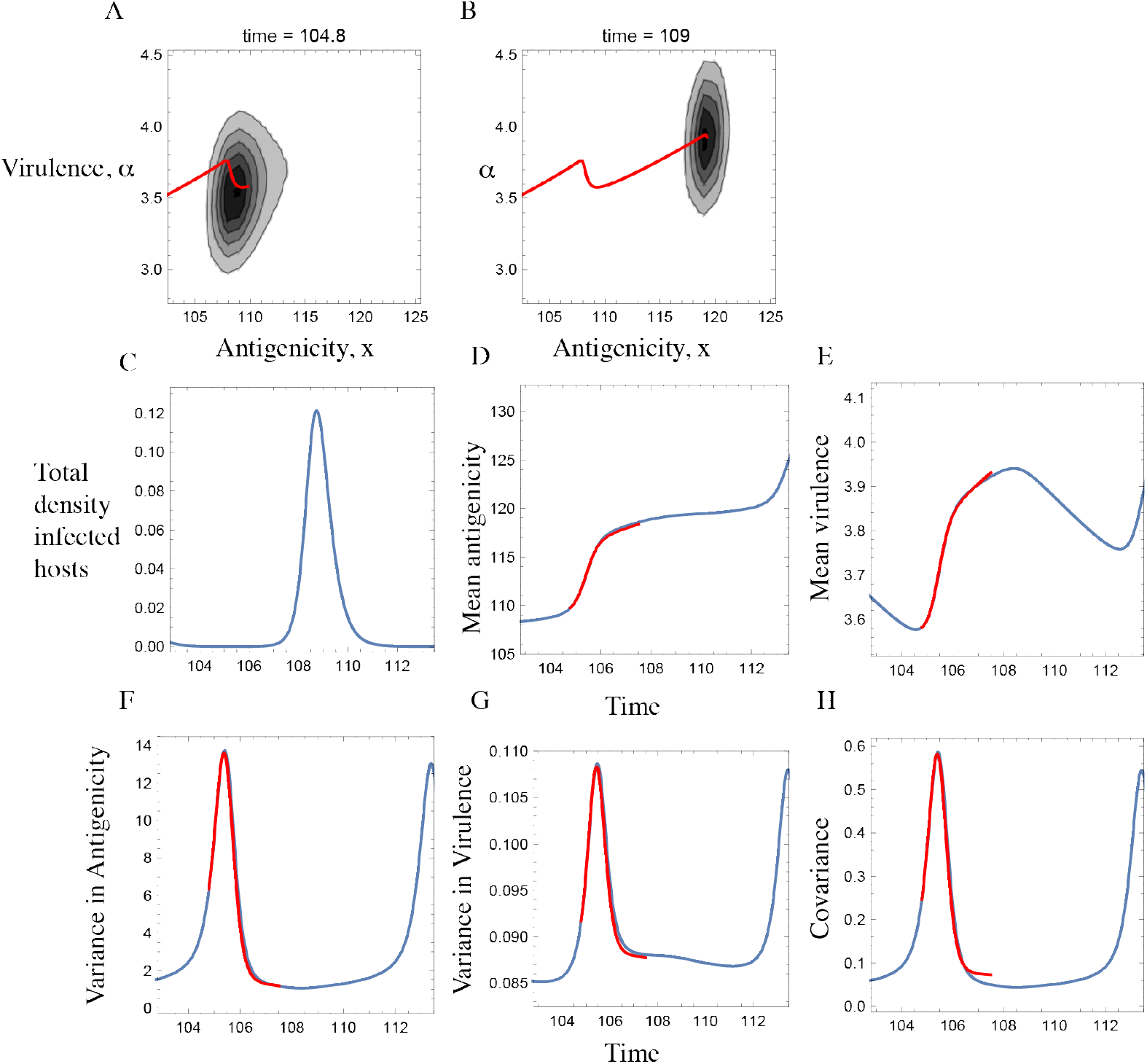
Oligomorphic dynamics prediction of the emergence of next strain in antigenicity-virulence coevolution. Panels (a) and (b) show the contour plots for the joint trait distribution observed in the same simulations at *t_s_* = 104.8 (at which the susceptibility distribution *s*(*x*) and initial moments in OMD are defined as explained in the main body) and *t_e_* = 109 (at which the OMD projection is terminated as the infected density increases around this time and then susceptibility distribution *s*(*x*) measured at *t* = *t_s_* no longer stays constant). The overlaid dotted curves are the trajectory of mean traits 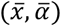 up *t* = *t_s_* and *t* = *t_e_* observed in the simulation. Panels (c), (d) and (e) show the dynamics of the total density of infected hosts, mean antigenicity and mean virulence, respectively. Parameters: *γ* = 0.5, 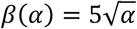, *D_x_* = 0.005, *D_α_* = 0.0002. As in Fig 2 we assume a Gaussian cross-immunity kernel, *σ*(*x* – *y*) = exp(−(*x* – *y*)^2^/2*ω*^2^), with width *ω* = 5. The oligomorphic dynamics describing the changes in the frequency *p*_0_(*t*) = 1 – *p*_1_(*t*) of the currently prevailing morph at time *t_s_* and the frequency *p*_1_(*t*) of upcoming morph, the mean antigenicity 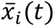 and mean virulence 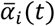 of the two morphs (*i* = 0,1), and the within-morph variances 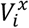 and 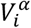 in antigenicity and virulence, as well as the within-morph covariance *C_i_*(*t*) between antigenicity and virulence in each morph (*i* = 0,1) are defined as (A44)–(A49) in Appendix S2.

## Discussion

We have shown how antigenic escape selects for more acute, virulent infectious diseases with higher transmission rates that cause increased mortality in infected hosts. This result is important given the number of important infectious diseases such as seasonal influenza that have epidemiology driven by antigenic escape. Until recently the evolution of virulence literature has mostly focused on equilibrium solutions that in simple models lead to the classic idea that pathogens evolve to maximize their basic reproductive number R_0_ [8,10–14]. Our results show that the process of antigenic escape leading to the continual replacement of strains [23–26], creates a dynamical invasion process that in and of itself selects for more acute, fast transmitting, highly virulent strains that do not maximize R_0_. This has parallels with the finding that more acute strains are selected temporally at the start epidemics [24–26], but critically, in our case the result is not a short-term transient outcome. Rather, the eco-evolutionary process leads to the long-term persistence of more acute strains with higher transmission rates causing higher virulence. As such, antigenic escape may be an important driver of high virulence in infectious disease.

In the simpler case where there is no cross immunity, there is a travelling wave of new strains invading due to antigenic escape. In this case we can use established methods to gain analytical results that not only predict the speed of change of the strains, but also the evolutionarily stable virulence. With our model’s assumptions, without antigenic escape we would get the classic result of the maximization of the reproductive number R_0_ [8,10–14], but once there is antigenic escape we show analytically that the intrinsic growth rate of the infectious disease *r* is maximized. Maximizing the intrinsic growth rate leads to selection for higher transmission leading to a much higher virulence. Effectively this is the equivalent of an infectious disease “live fast, die young” strategy. The outcome is due to the dynamical replacement of strains, with new strains invading the population continually leading to a continual selection for the strains that invade better [24–26]. As such we predict that any degree of antigenic escape will in general select for more acute faster transmitting strains with higher virulence in the presence of a transmission-virulence tradeoff.

Partial cross-immunity leads to a series of jumps in antigenic space that are characteristic of the epidemiology of a number of diseases and in particular, in humans, the well-known dynamics of influenza A & B [23–26]. Here a cloud of strains remains in antigenic space until there is a jump that, on average, overcomes the cross immunity and leads to the invasion of a new set of strains that are distant enough to escape the immunity of the resident strains [23–26]. In order to examine the evolutionary outcome in this scenario we applied a novel oligomorphic analysis [28]. Again, we find that antigenic escape selects for higher virulence with more acute faster transmitting strains being favored with again selection leading toward the maximization of the intrinsic growth rate *r*. Both our analysis and simulations show that in the long term the virulence increases until it reaches a new optima potentially of an order of magnitude higher than would be expected by the classic prediction of maximizing R_0_. We show that virulence increases between each antigenic jump, falling slightly at the next jump before increasing again until it reaches this new equilibrium. It is also important to note that diversity in both antigenicity and virulence increases as we move towards the next epidemic, maximizing just at the point when the jump occurs. This increase in diversity could in principle be used as a predictor of the next jump in antigenic space.

Clearly the virulence of any particular infectious disease depends on multiple factors, including both host and parasite traits, and critically the relationship between transmission and virulence. This makes comparisons of the virulence across different infectious diseases problematic since the specific trade-off relationship between transmission and virulence is often unknown. However, our model shows that antigenic escape will, all things being equal, be a driver of higher virulence favoring more acute strains. It is also important to note that since antigenic escape is a very general mechanism that selects for higher virulence it follows that we may see high virulence in parasites even when the costs in terms of reduced infectious period are substantial. Amongst the influenzas, influenza C does not show obvious antigenic escape [33] and is typically much less virulent than the other influenzas which are the classic examples of infectious disease with antigenic escape. Furthermore, Influenza A shows much more antigenic escape than Influenza B and again in line with our predictions typically influenza A is the more virulent [34,35]. Clearly these differences can be ascribed to multiple factors and indeed the higher virulence of influenza A is often posited to be due to a more recent zoonotic emergence [36]. However, our models suggest that the differences in the degree to which they show antigenic escape *per se* may also contribute to these differences. Clearly there are also highly virulent pathogens that do not show antigenic escape and a formal comparative analysis is confounded by multiple factors, but the evidence from influenza is consistent with our predictions.

An important implication of our work is that it follows that antigenic escape selects for strains with a higher virulence than the value that maximizes R_0_ and therefore leads to the evolution of infectious diseases with lower R_0_. One simplistic implication of this is that, from this point of view, diseases with antigenic escape may be easier to eliminate and control with vaccination. Of course, in practice the opposite is often true since producing an effective vaccine is much more problematic when there is antigenic escape [33,37]. In addition, it follows that epidemics will tend to be less explosive than they otherwise would have been, having a lower peak but lasting longer, with evolution here effectively ‘flattening the curve’. Infectious diseases that show jumping antigenic escape are characterized by repeated epidemics, but our work suggests that due to the selection for a lower R_0_ the eco-evolutionary feedback will have significantly impacted the pattern of these epidemics. This is an important example of a dynamical ecological/epidemiology feedback into evolutionary outcomes that in turn then feedbacks into the epidemiology characteristics of the disease.

We have used oligomorphic dynamics [28] to make predictions on the waiting times and outcomes of the antigenic jumps in our model with cross immunity. This approach tracks both mean trait values and the changes in trait variances in models with explicit ecological dynamics. As such it combines aspects of eco-evolutionary theory [38,39] and quantitative genetics approaches [40,41] to provide a more complete understanding of the evolution of quantitative traits. Our approach can assume a wide range of different ecological and evolutionary time scales and therefore allows us to address fundamental questions on eco-evolutionary feedbacks and on the separation between ecological and evolutionary time scales. This is important since it allows us to test the implications of the different assumptions of classical evolutionary theory and to better understand the role of eco-evolutionary feedbacks on evolutionary outcomes. Furthermore, the approach can be applied widely to model transient dynamics, and to predict the waiting times and extent of diversification that occurs in a range of contexts [28,42]. Here it allowed us to predict evolutionary outcomes analytically in the dynamical context of antigenic escape.

Broadly, our results emphasize that epidemiological dynamics may have important implications to the evolution of infectious disease. Here repeated invasions driven by antigenic escape mutants were shown to lead to the long term evolution of more acute virulent pathogens. It has previously been shown that newly invading parasites will temporarily not be at the classic R_0_ optima [19,32], but our model shows that this more acute parasite state is a long term optima when there is antigenic escape. Furthermore, this long term optima leads to a more acute severe disease which is clearly of particular importance in human health. Human coronaviruses can evolve antigenically to escape antibody immunity [43] and moreover in principle epidemics of new variants as a novel disease adapts to the host will lead to equivalent dynamics and evolutionary outcomes. It also follows that interventions that impact epidemiological dynamics may also have impacts on the evolution of pathogen traits such as virulence or transmission and it is therefore critical that we have a broader theory of the evolution of virulence that predicts the implications of these epidemiological effects. Furthermore, these dynamical feedbacks are likely to be important in a range of contexts beyond infectious disease and the analytical approaches we use here are likely to be useful in understanding dynamical evolutionary outcomes.

## Acknowledgements

This study was supported by grants R01 GM122061-03 and NSF-DEB-2011109 to MB, ANR JCJC grant ANR-16-CE35-0012-01 to SL and the ESB Cooperation Program, The Graduate University for Advanced Studies, SOKENDAI.

## Appendix S1: Oligomorphic dynamics of antigenic escape

We consider a model of antigenic escape of a pathogen from host herd immunity on a onedimensional antigenicity space (*x*), which describes the changes in the density *S*(*t, x*) of hosts that are susceptible to antigenicity strain *x* of pathogen at time *t*, and the density *I*(*t, x*) of hosts that are currently infected and infectious with antigenicity strain *x* of pathogen at time *t*:

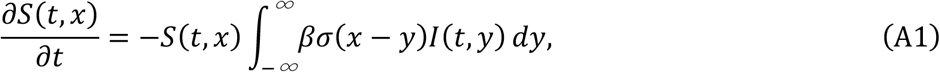

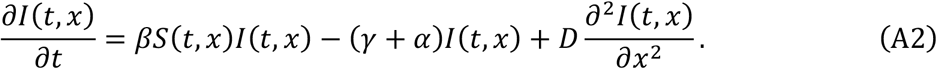

where *β, α*, and *γ* are the transmission rate, virulence (additional mortality due to infection), and recovery rate of pathogens which are independent of antigenicity. *σ*(*x* – *y*) denotes the degree of cross immunity: a host infected by strain *y* of pathogen acquires perfect cross immunity with probability *σ*(*x* – *y*) but fails to acquire any cross immunity with probability 1 – *σ*(*x* – *y*) (this is called polarized cross immunity by Gog and Grenfell 2002). The cross-immunity kernel *σ*(*x* – *y*) is assumed to be a decreasing function of the distance |*x* – *y*| between strains *x* and *y*. When a new strain with antigenicity *x* = 0 is introduced at time *t* = 0, the initial host population is assumed to be susceptible to any antigenicity strain of pathogen: *S*(0, *x*) = 1. 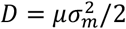 is one half of the mutation variance for the change in antigenicity, representing random mutation in the continuous antigenic space.

### Susceptibility profile molded by the primary outbreak

We first analyze the dynamics of the primary outbreak of a pathogen and derive the resulting susceptibility profile, which can be viewed as the fitness landscape subsequently experienced by the pathogen. For simplicity we assume that mutation can be ignored during the first epidemic started with the antigenicity stain *x* = 0. The density *S*_0_(*t*) = *S*(*t*, 0) of hosts that are susceptible to the currently prevailing antigenicity strain *x* = 0, as well as the density *I*_0_(*t*) = *I*(*t*, 0) of hosts that are currently infected by the focal strain change with time as

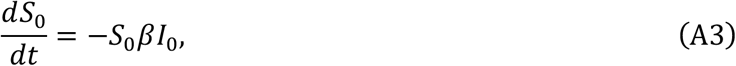

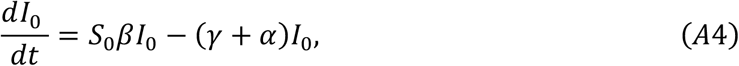

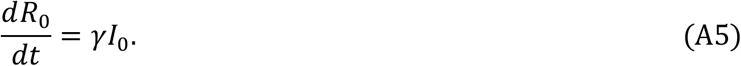

with *S*_0_(0) = 1, *I*_0_(0) ≈ 0, and *R*_0_(0) = 0. The final size of the primary outbreak,

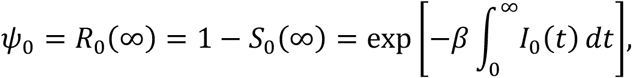

is determined as the unique positive root of

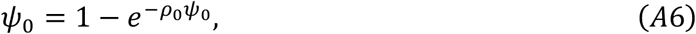

where *ρ*_0_ = *β*/(*γ* + *α*) > 1 is the basic reproductive number [Anderson and May, 1991]. Associated with this epidemiological change, the susceptibility profile *S_x_*(*t*) = *S*(*t, x*) against antigenicity *x* (*x* ≠ 0) other than the currently circulating strain (*x* = 0) changes by cross immunity as

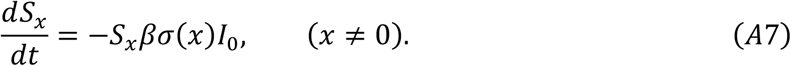

Integrating both sides of (A7) from *t* = 0 to *t* = ∞, we see that the susceptibility profile *s*(*x*) = *S_x_*(∞) after the primary outbreak at *x* = 0 is

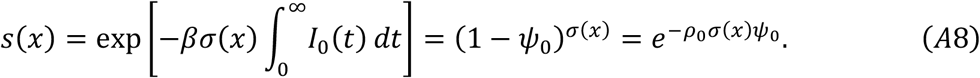

where the last equality follows from (A6). The susceptibility can be effectively reduced by cross-immunity when the primary strain has a large impact (the fraction of hosts remained uninfected, 1 − *ψ*_0_, is small) and when the degree of cross immunity is strong (*σ*(*x*) is close to 1). With a strain antigenically very close to the primary strain (*x* ≈ 0), the cross immunity is very strong (*σ*(*x*) ≈ 1) so that the susceptibility against strain *x* is nearly maximally reduced *s*(*x*) ≈ 1 – *ψ*_0_. With a strain antigenically distant from the primary strain, *σ*(*x*) becomes substantially smaller than 1, making the host more susceptible to the strain. For example, if the cross immunity is halved (*σ*(*x*) = 0.5) from its maximum value 1, then the susceptibility to that strain is as large as (1 – *ψ*_0_)^0.5^. If a strain is antigenically very distant from the primary strain, *σ*(*x*) ≈ 0, and the host is nearly fully susceptibility to the strain (*s*(*x*) ≈ 1).

**Figure S1:**
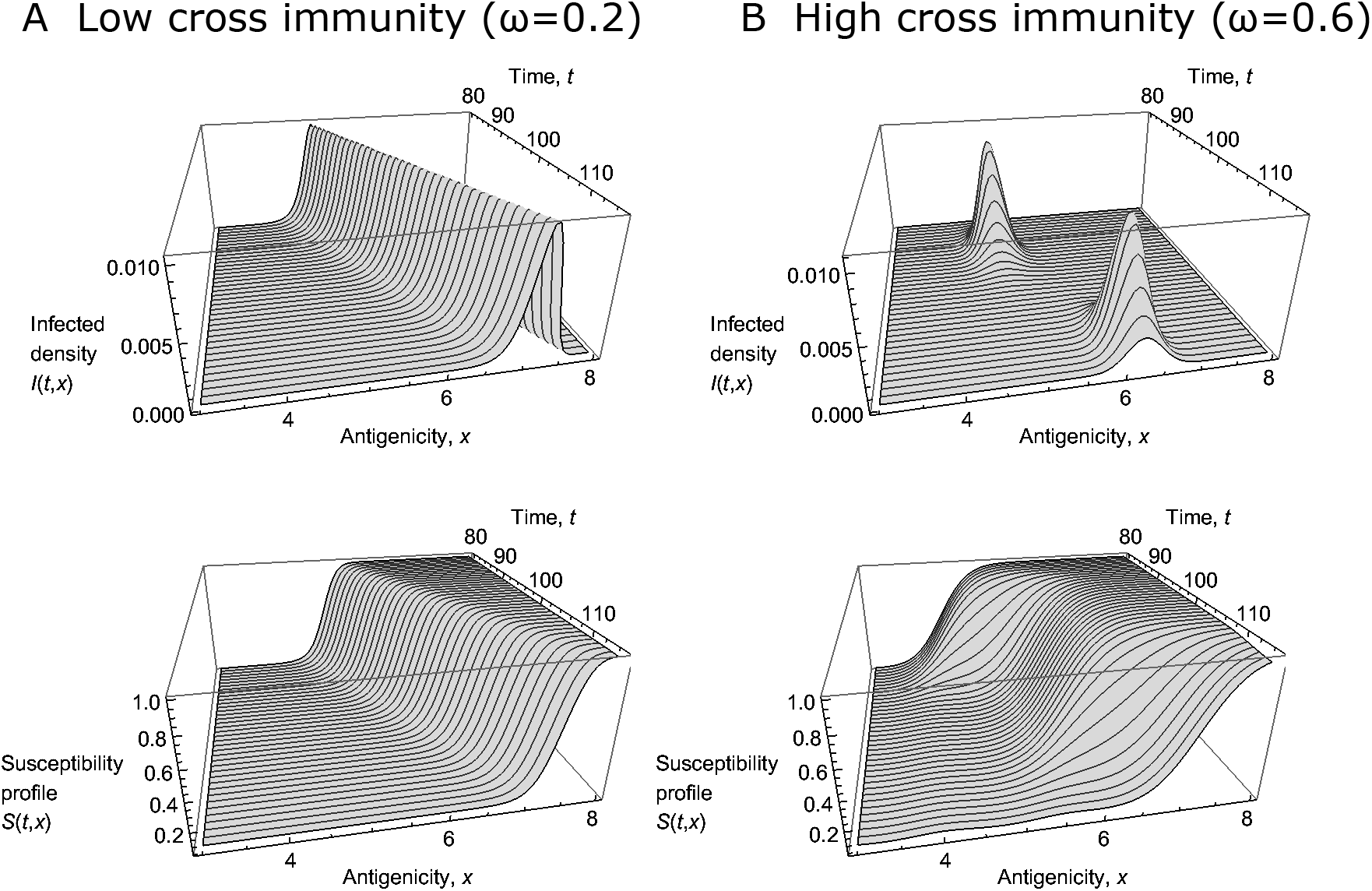
Continuous antigenic drift (A) and periodical antigenic shits (B) of the model. The orange colored surface denotes the infected density *I*(*t, x*) varying in time *t* and intigenicty *x*, and the yellow colored surface denotes the density of hosts *S*(*t, x*) that are susceptible to antigenicity strain *x* of pathogen at time *t*. The width of cross immunity *ω* = 0.2 in (A) and *ω* = 0.6 in (B). A continuous antigenic drift solution (travelling wave of a fixed shaped profile with a constant wave speed) loses stability around *ω* = 0.4 and the departure from the travelling wave increases as *ω* increases (C). The wave speeds stay nearly constant and agree with (3) when *ω* is varied (D). Other parameters are *β* = 2, *u* = 0.001, *α* = 0.1, *γ* = 0.5, and *D* = 0.001.

### Threshold for antigenic escape

Of particular interest is the threshold antigenicity distance *x_c_* that allows the antigenic escape, i.e. any antigenicity strain more distant than this threshold from the primary strain (*x* > *x_c_*) can increase when introduced after the primary outbreak. Such a threshold is determined from

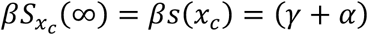

or

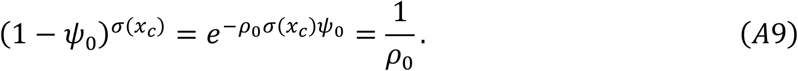

With a specific choice of cross-immunity profile,

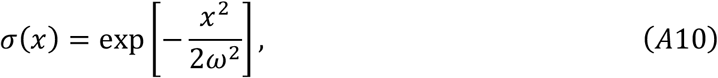

the threshold antigenicity beyond which the virus can increase in the susceptibility profile *s*(*x*) after the primary outbreak is, from

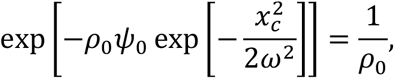

and taking logarithm of both sides twice,

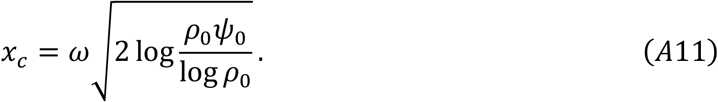

### Oligomorphic dynamics

Integrating both sides of (A2) over the whole space, we obtain the dynamics for the total density of infected hosts, 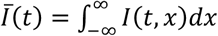:

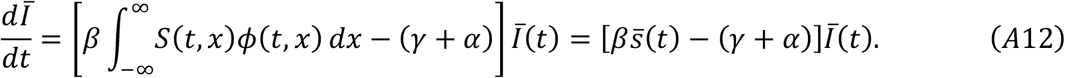

where

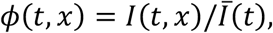

is the relative frequency of antigenicity strain *x* circulating at time *t*, and

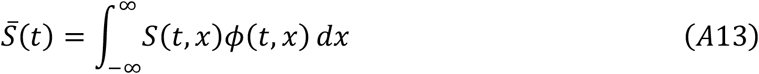

is the mean susceptibility experienced by currently circulating pathogens. The dynamics for the relative frequency *ϕ*(*t, x*) of pathogen antigenicity is

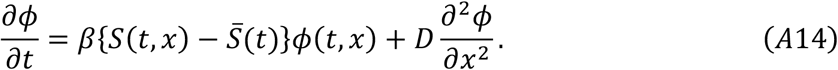

As in Sasaki and Dieckmann (2011), we decompose the frequency distribution to the sum of several morph distributions (oligomorphic decomposition) as

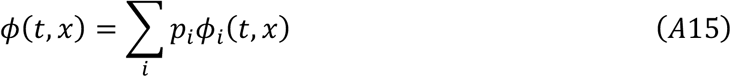

where *p_i_*(*t*) is the frequency of morph *i* and *ϕ_i_*(*t, x*) is within-morph distribution of antigenicity. By definition, Σ_*i*_ *p_i_* and 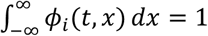. Let

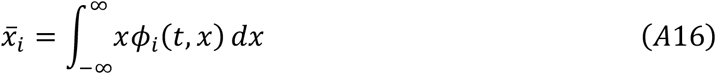

be the mean antigenicity of a morph and

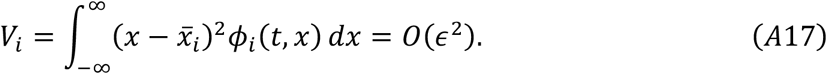

be the within-morph variance of each morph, which is assumed to be small, of the order of *ϵ*^2^. Let us denote the mean susceptibility of host population for viral morph *i* by 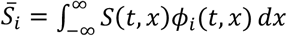. As shown in Sasaki and Dieckmann (2011), the dynamics for viral morph frequency is expressed as

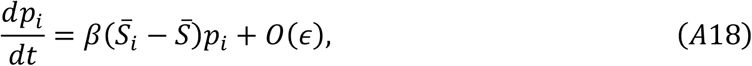

while the dynamics for the within-morph distribution of antigenicity is

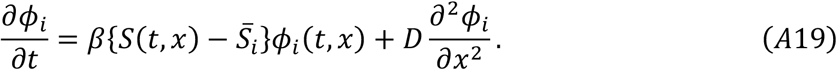

From this, the dynamics for the mean antigenicity of a morph,

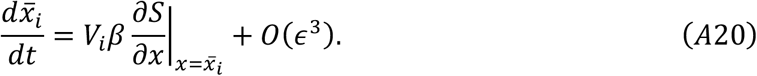

and the dynamics for the within-morph variance of a morph

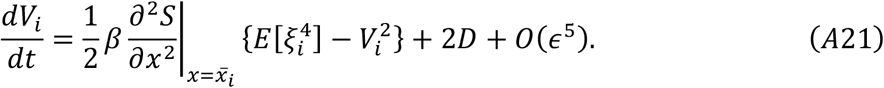

are derived, where 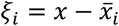 and 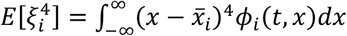 is the fourth central moment of antigenicity around the morph mean. Assuming that the morph distribution is and remains normal (Gaussian closure), 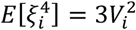 and hence Eq. (A21) becomes

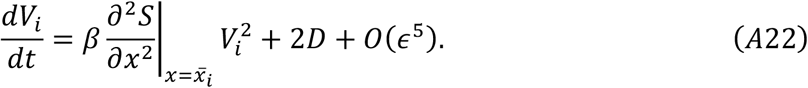

### Second outbreak predicted by OMD

Equations (A18), (A20) and (A22) are general, but they rely on a full knowledge of the dynamics of the susceptibility profile *S*(*t, x*). In order to make further progress, we use an additional approximation by substituting Eq. (A9), the susceptibility profile over viral antigenicity space after the primary outbreak at *x* = 0 and before the onset of the second outbreak at a distant position. We keep track of two morphs at positions *x*_0_(*t*) and *x*_1_(*t*), where the first morph is that caused the primary outbreak at *x* = 0, and the second morph is that emerged in the range *x* > *x_c_* beyond the threshold antigenicity *x_c_* defined in Eq. (A9) (and (A11) for a specific form of *σ*(*x*)) as the source of the next outbreak.

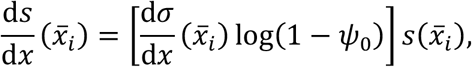

and

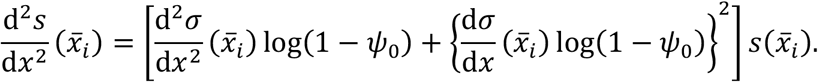

Therefore, the frequency, mean antigenicity, and variance of antigenicity of an emerging morph (*i* = 1) change respectively as

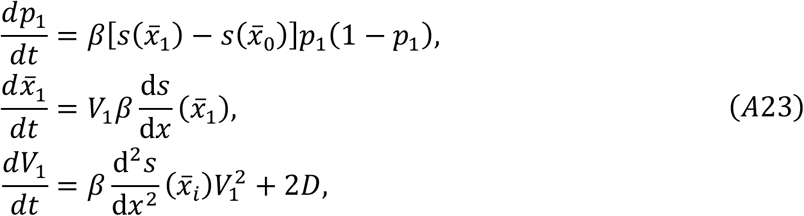

The predicted change in the mean antigenicity is plotted by integrating Eq. (A23). As initial condition, we choose the time when a seed of second peak in the range *x* > *x_c_* first appears, and then compute the mean trait as

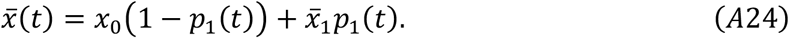

In the case of Figure 1, where *β* = 2, *γ* + *α* = 0.6, *D* = 0.001, and *ω* = 2, the final size of epidemic for the primary outbreak, defined as (7) was *ψ* = 0.959, and the critical antigenic distance for the increase of pathogen strain obtained from (A22) was *x_c_* = 2.795. The initial condition for the oligomorphic dynamics (A23) for the second morph was then *p*_1_(*t*_0_) = 1.6 x 10^−8^, 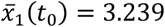, *V*_1_(*t*_0_) = 0.2675 at *t*_0_ = 41. In Figure 1, the predicted trajectory for the mean antigenicity (A24) is plotted as red curve, together with the mean antigenicity change observed in simulation (blue curve).

### The accuracy of predicting with OMD the antigenicity and the timing of the second outbreak

Here we describe how the initial conditions for oligomorphic dynamics, i.e., the frequency, the mean antigenicity and the variance in antigenicity of the morph that caused the primary outbreak and the morph which may cause the second outbreak are determined. We then show how the accuracy in prediction of the second outbreak depends on the timing for prediction.

We divide the antigenicity space into two at *x* = *x_c_* above which the pathogen can increase under the given susceptibility profile after the primary outbreak, but below which the pathogen cannot increase. We then take relative frequencies of pathogens above *x_c_* and below *x_c_*, and the conditional mean and variance in these separated regions to set the initial frequencies, means, and variances of the morphs at the time *t*_0_ when we predict the second outbreak:

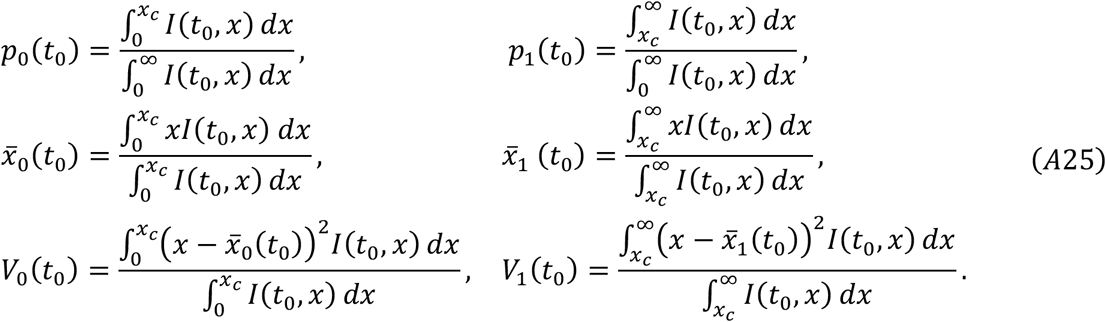

We then compare the trajectory for mean antigenicity change observed in simulation (blue curve in Figure 1) and the predicted trajectory (red curve in Figure 1) for mean antigenicity (A24) by integrating oligomorphic dynamics (A23) with initial condition (A25) at time *t* = *t*_0_. Figure S1 shows how the accuracy of prediction, measured by the Kullback-Leibler divergence or L2 norm between these two trajectories depends on the timing chosen for the prediction, *t*_0_. The second outbreak occurs around t = 54.6 where mean antigenicity jumps from around 0 to around 5. The prediction with OMD is accurate if it is made for *t*_0_ > 40. Figure 1 is drawn for *t*_0_ = 41 where the second peak is about to emerge (see Figure S2). Even for the latest prediction for *t*_0_ = 51 in Figure S2, the morph frequency of the emerging second quasispecies was only 0.3%, so the prediction is still worthwhile to make.

Figure S2 shows that the prediction power is roughly constant (albeit with a wiggle) for 5 < *t*_0_ < 30 (the predicted timings are 10-15% longer than actual timing for 5 < *t*_0_ < 30), and steadily improved for *t*_0_ > 30. When the prediction is made very early <(*t*_0_ < 5) the deviations are larger.

**Figure S2.**
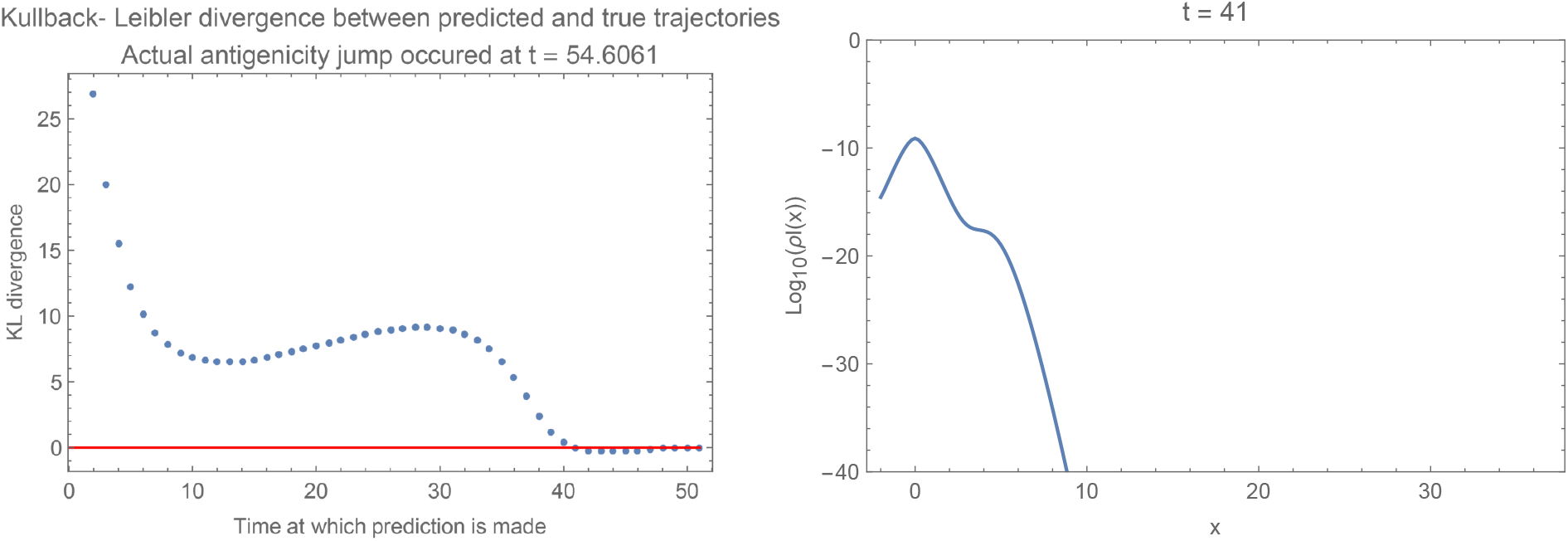
The accuracy of prediction of the second outbreak as a function of the prediction timing, using either the Kullback-Leibler divergence (left panel) (A) The KL divergence plotted here is 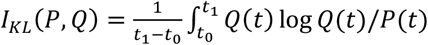, where *Q*(*t*) is the trajectory for mean antigenicity observed in simulation, and *P*(*t*) is the corresponding trajectory obtained with OMD with using the data at *t* = *t*_0_ being used to set the initial condition. The right end of the comparison in time horizon is set to *t*_1_ = 80 when the second outbreak is over. (B) The logarithmic density of antigenicity strains, *I*(*t, x*), at *t* = 41. A tiny seed for the second morph around *x* = 6 is visible.

## Appendix S2: Oligomorphic dynamics for the joint evolution of antigenicity and virulence

Let *s*(*x*) be the susceptibility of the host population against antigenicity *x* a specific susceptibility profile is given by (A8) in Appendix S1 with cross-immunity function *σ*(*x*) and the final size *ψ*_0_ of epidemic of the primary outbreak. Note that as in appendix S1, the susceptibility profile is in general a function of time. The density *I*(*x, α*) of hosts infected by a pathogen of antigenicity *x* and virulence *α* changes with time, when rare, as

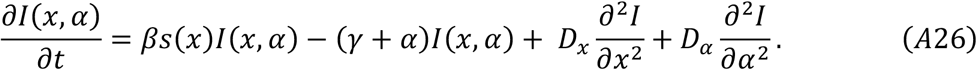

The change in the frequency 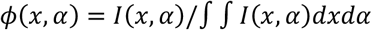 of a pathogen with antigenicity *x* and virulence *α* follows

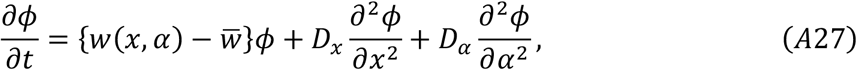

where

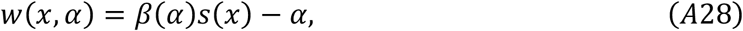

is the fitness of a pathogen with antigenicity *x* and virulence *α* and 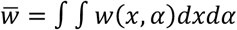 is the mean fitness.

Let us decompose the joint frequency distribution *ϕ*(*x, α*) of the viral quasispecies as (oligomorphic decomposition):

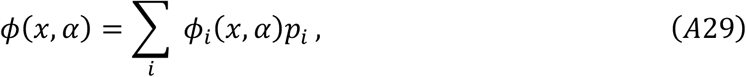

where *ϕ_i_*(*x, α*) is the joint frequency distribution of antigenicity *x* and virulence *α* in morph *i* 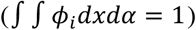 and *p_i_* is the relative frequency of morph *i*(Σ_*i*_ *p_i_* = 1). The frequency of morph *i* then changes as

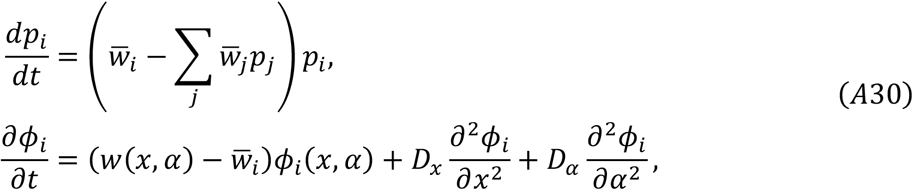

where 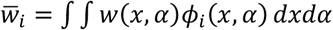 is the mean fitness of morph *i*.

Assuming that the traits are distributed narrowly around the morph means 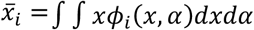 and 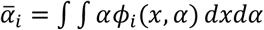, so that 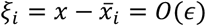 and 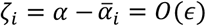 where *ϵ* is a small constant, we expand the fitness *w*(*x, α*) around the means 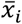 and 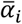 of morph *i*,

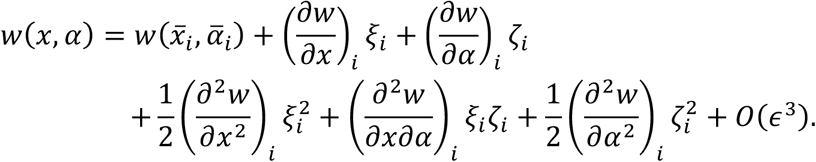

Substituting this and

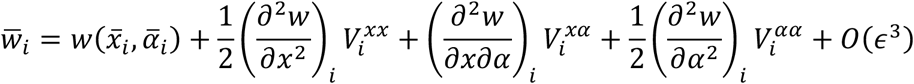

into (A30), we have

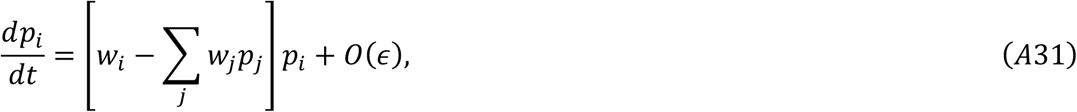

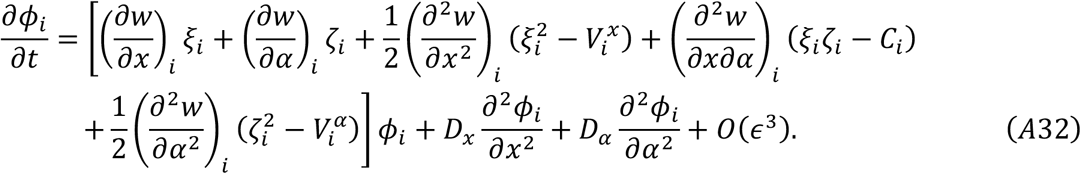

where 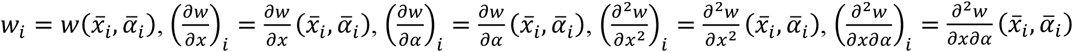, and 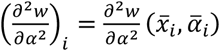 are fitness and its first and second derivatives evaluated at the mean traits of morph *i*, and

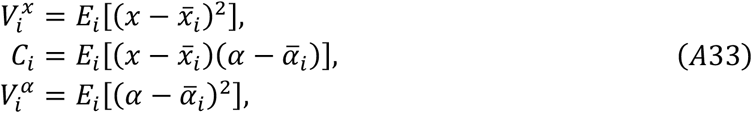

are within-morph variances and covariance of the traits of morph *i*. Here, 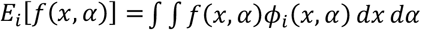 denotes taking expectation of a function *f* with respect to the joint trait distribution *ϕ_i_*(*x, α*) of morph *i*.

Substituting (A32) into the change in the mean antigenicity of morph *i* we have

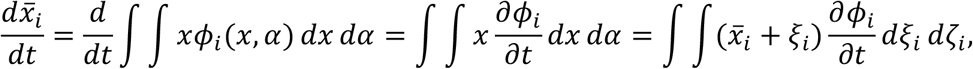

we have

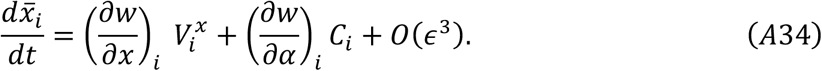

Similarly, the change in the mean virulence of morph *i* is expressed as

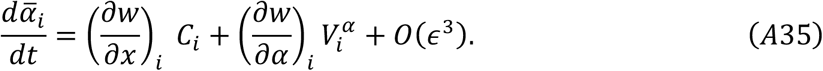

Equations (A34)–(A35) from the mean trait change is summarized in a matrix form as

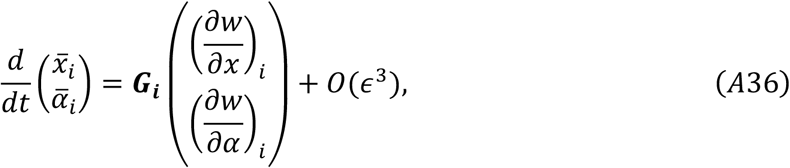

where

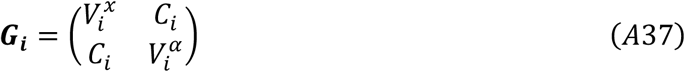

is the variance-covariance matrix of the morph *i*.

Substituting (A32) into the right-hand side of the change in variance of antigenicity of morph *i*,

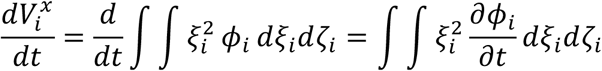

and those in the change in the other variance and covariance, we have

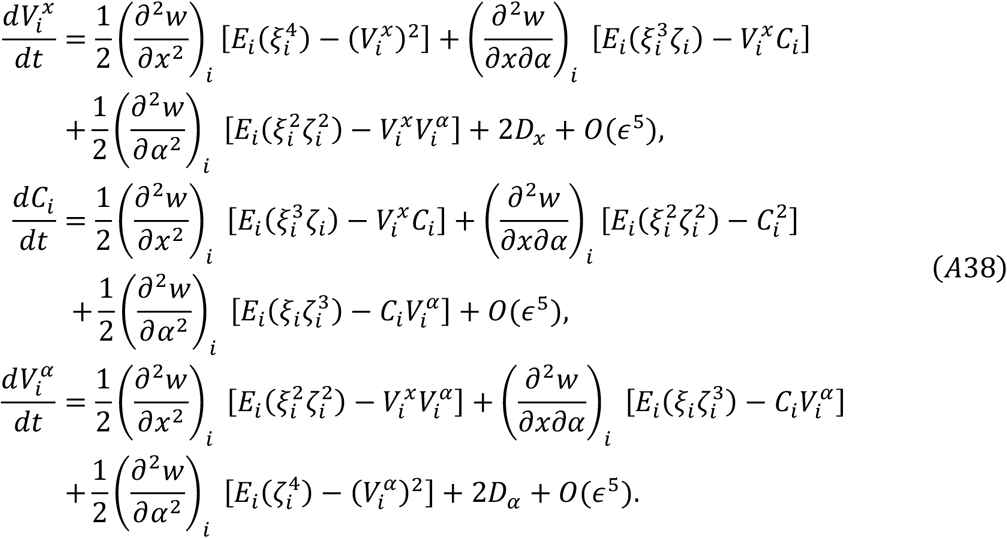

If we assume that antigenicity and virulence within a morph follow two-dimensional Gaussian distribution for given means, variances and covariance, we should have 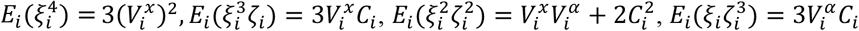, and 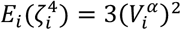, and hence

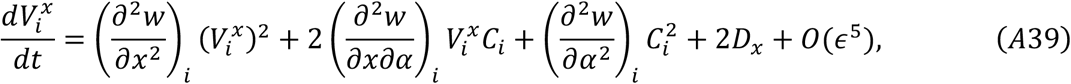

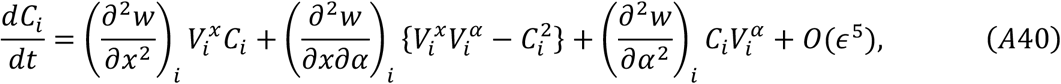

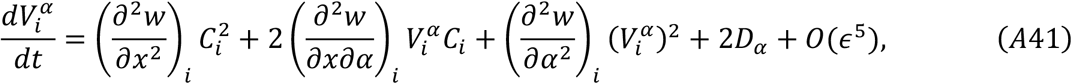

Eqs. (A39)–(A41) are rewritten in a matrix form as

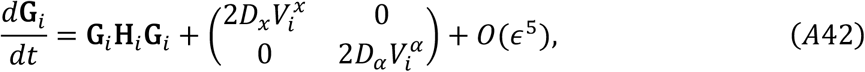

where

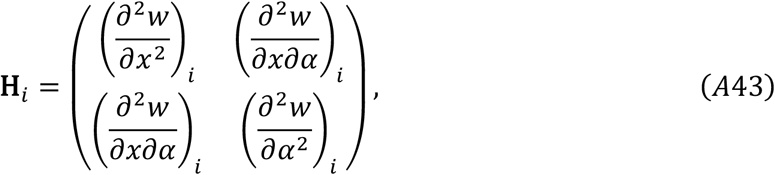

is the Hessian of the fitness function of the morph *i*.

In our case (A26) of the joint evolution of antigenicity and virulence of a pathogen after its primary outbreak, the fitness function is given by *w*(*x, α*) = *β*(*α*)*s*(*x*) – *α*, and hence 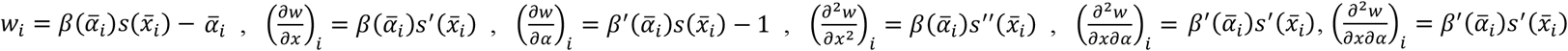, and 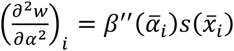, where a prime on *β*(*α*) and *s*(*x*) denotes differentiation by *α* and *x*, respectively. Substituting these into the dynamics for morph frequencies (A31), for morph means (A34)–(A35), and for within-morph variance and covariance (A39)–(A41), we have

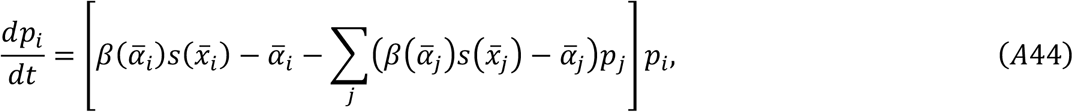

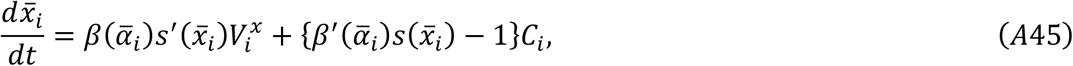

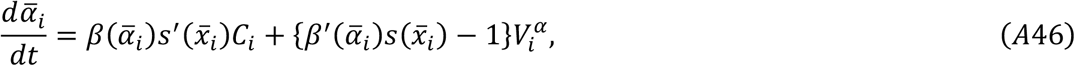

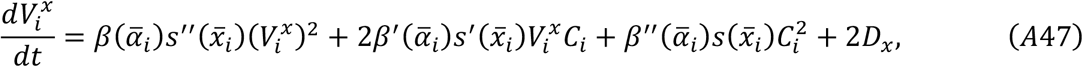

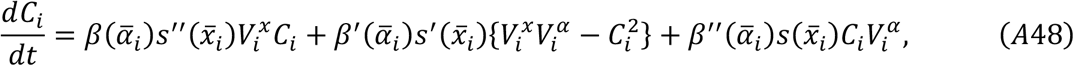

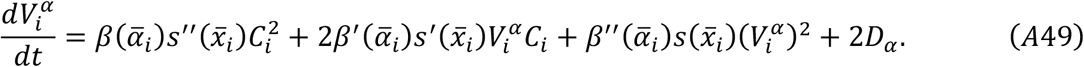

Equations (A44)–(A49) describe the oligomorphic dynamics of the joint evolution of antigenicity and virulence of a pathogen for a given host susceptibility profile *s*(*x*) over pathogen antigenicity. Of particular interest is whether the antigenicity evolution or virulence evolution accelerates each other by allowing them evolving simultaneously than when they evolve alone. After the primary outbreak at a given antigenicity, say *x* = 0, the susceptibility *s*(*x*) of host population increases due to cross-immunity as the distance *x* > 0 from the antigenicity at the primary outbreak. Combining this with the positive tradeoff between transmission rate and virulence, we see that 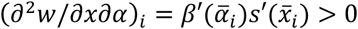, and then from (A48) we see that the within-morph covariance between antigenicity and virulence stays positive if its initial value is nonnegative:

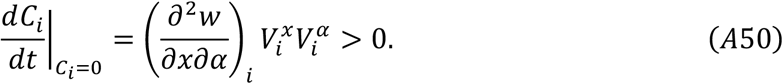

If all second moments are sufficiently small initially for an emerging morph, a quick look at the linearization of (A47)–(A49) around 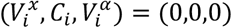 indicates that both 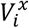 and 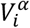 become positive due to the random generation of variance by mutation, *D_x_* > 0 and *D_α_* > 0, while the covariance stays close to zero. Then (A50) guarantees that first move of covariance is from zero to positive, which then guarantees that *C_i_* > 0 for all *t*. Therefore, the second term in (A34) is positive until the mean virulence reaches its optimum (*β*′(*α*)*s*(*x*) = 1). This means that joint evolution with virulence accelerates the evolution of antigenicity. The same is true for virulence evolution: the first term in (A35) (which denotes the associated change in virulence due to the selection in antigenicity through genetic covariance between them) is positive, indicating that joint evolution with antigenicity accelerates the virulence evolution.

### Numerical example

Figure 5 in the main body shows the oligomorphic dynamics prediction of the emergence of next strain in antigenicity-virulence coevolution. In order to make progress numerically we assume *s*(*x*) to be constant in the following analysis because we are interested in the process between the end of the primary outbreak and the emergence of the next antigenicity virulence morph. A The partial differential equations for the density of host *S*(*t, x*) susceptible to the antigenicity strain *x* at time *t*, and the density of hosts infected by pathogen strain with antigenicity *x* and virulence *α* are

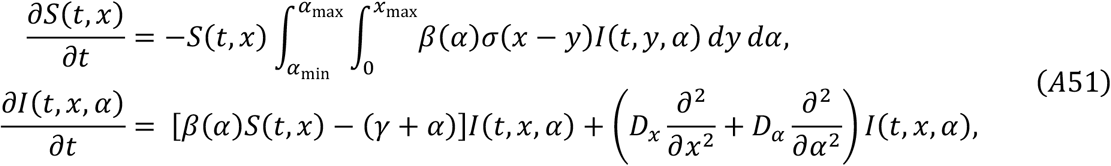

with the boundary conditions (*∂S*/*∂x*)(*t*, 0) = (*∂S*/*∂x*)(*t, x*_max_) = 0, (*∂I/∂x*)(*t*, 0, *α*) = (*∂I*/ *∂x*)(*t, x*_max_, 0) = 0, (*∂I/∂x*)(*t, x, α*_min_) = (*∂I/∂x*)(*t, x, α*_max_) = 0, and the initial conditions *S*(0, *x*) = 1, and *I*(0, *x*, *α*) = *ϵδ*(*x*)*δ*(*α*) where *δ*(·) is delta function and *ϵ* = 0.01. The trait space is restricted in a rectangular region: 0 < *x* < *x*_max_ = 300 and *α*_min_ = 0.025 < *α* < 10 = *α*_max_. Oligomorphic dynamics prediction for the joint evolution of antigenicity and virulence is applied for the next outbreak after the outbreak with the mean antigenicity around *x* = 108 at time *t* = 102. The susceptibility of host to antigenicity strain *x* at *t*_0_ = 104.8 after the previous outbreak peaked around time *t* = 102 came to an end is

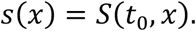

This susceptibility profile remains unchanged until the next outbreak starts, and hence the fitness of a pathogen strain with antigenicity *x* and virulence *α* is given by

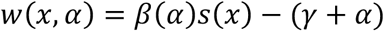

We divide the pathogen quasispecies into two morphs at time *t*_0_ at the threshold antigenicity *x_c_* above which the net growth rate of pathogen strain under the given susceptibility profile *s*(*x*) and the mean antigenicity becomes positive:

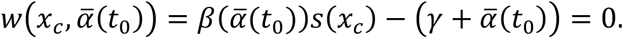

The initial frequency and the moments of two morphs, the strain 0 with *x* < *x_c_* and the strain 1 with *x* > *x_c_* are then calculated respectively from the joint distribution *I*(*t*_0_, *x, α*) in the restricted region {(*x, α*); 0 < *x* < *x_c_*, *α*_min_ < *α* < *α*_max_} and that in the restricted region {(*x, α*); *x_c_* < *x* < *x*_max_, *α*_min_ < *α* < *α*_max_}. The frequency *p*_1_ of the morph 1 (the frequency of the morph 0 is given by *p*_0_ = 1 – *p*_1_), the mean antigenicity 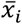 and mean virulence 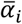 of the morph *i*, and variances and covariance, 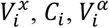 of the morph *i* (*i* = 0,1) follow (A44)–(A49), where the dynamics for the morph frequency (A44) is simplified in this two morph situation as

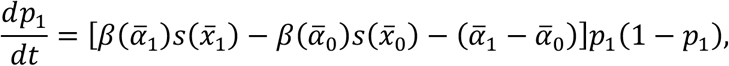

with *p*_0_(*t*) = 1 – *p*_1_(*t*). This is iterated from *t* = *t*_0_ = 104.8 to *t_e_* = 107. The frequency *p*_1_ of the new morph, the population mean antigenicity 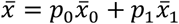, virulence 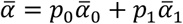, variance in antigenicity 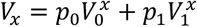, covariance between antigenicity and virulence *C* = *p*_0_*C*_0_ + *p*_1_ *C*_1_, and variance in virulence 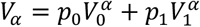 are overlayed by red thick curves on the trajectories of moments observed in full dynamics (A51).

In the panel (a) of Figure 5, the dashed vertical line represents the threshold antigenicity *x_c_* above which 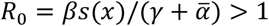 at *t* = *t_s_* = 104.8 where oligomorphic dynamics (OMD) prediction is attempted. Two morphs are then defined according to whether or not the antigenicity exceeds a threshold *x* = *x_c_*: the resident morph (morph 1) is represented as the dense cloud to the left of *x* = *x_c_* and the second morph (morph 2) consisting of all the genotypes to the right of *x* = *x_c_* with their *R*_0_ greater than one. The within-morph means and variances are then calculated in each region. The relative total densities of infected hosts in the left and right regions defines the initial frequency of two morphs in OMD. A 2D Gaussian distribution is assumed for within-morph trait distributions to have the closed moment equations as explained before. Using these as the initial means, variances, covariances of the two morphs at *t* = *t_s_*. The oligomorphic dynamics for 11 variables (relative frequency of morph 1, mean antigenicity, mean virulence, variances in antigenicity and virulence and their covariance in morph 0 and 1) is integrated up to *t* = *t_e_*. The results are shown in red curves in the panels of second and third rows, which are compared with the simulation results (blue curves).

The panels (c)-(e) in Figure 5 show the change in total infected density (c), mean antigenicity (d), and mean virulence (e). Red curves show the prediction by oligomorphic dynamics from the initial moments of each morph at *t* = *t_s_* to the susceptibility distribution *s*(*x*) = *S*(*t_s_, x*). Red curves in the (d) and (e) show the OMD prediction, which is compared with the simulation results (blue curves). The OMD predicted mean antigenicity, for example, is defined as

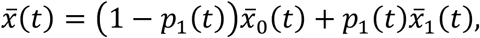

where *p*_1_(*t*) is the frequency of morph 1, 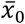 and 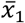 are the mean antigenicity of morph 0 and 1.

The red curves in the third row of Figure 5 show the OMD-predicted changes in the variance in antigenicity, variance in virulence, and covariance between antigenicity and virulence, which are compared with the simulation results (blue curves). The OMD predicted covariance, for example, is defined as

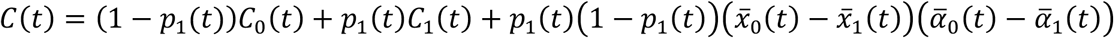

where *C*_0_(*t*) and *C*_1_(*t*) are antigenicity-virulence covariance in morph 0 and 1, and 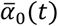 and 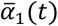 are mean virulence of morph 0 and 1.

## Appendix S3: Selection for maximum growth rate

In this appendix, we show that a pathogen that has the strategy maximizing the growth rate in a fully susceptible population is evolutionarily stable in the presence of antigenic escape.

At stationarity, the travelling wave profiles of 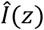 and 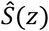 along the moving coordinate, *z* = *x* – *νt*, that drifts constantly to right with the speed *ν* are defined as

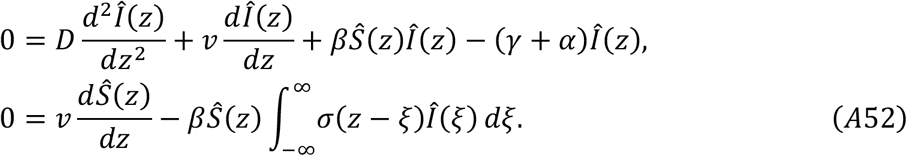

with 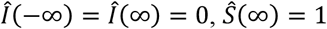.

Let *j*(*t, x*) be the density of a mutant pathogen strain with virulence *α*′ and transmission rate *β*′ that is introduced in the host population where the resident strain is already established (A46). For the initial transient phase in which he density of mutant is sufficiently small, we have an equation for the change in *J*(*t, z*) = *j*(*t, x*):

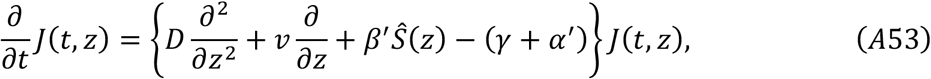

with the initial condition *J*(0, *z*) = *ϵδ*(*z*), where *ϵ* is a small constant and *δ*(·) is Dirac function.

Consider a system

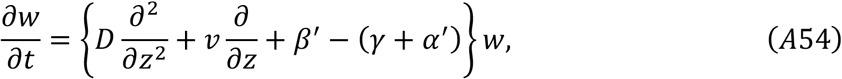

with *w*(0, *z*) = *J*(0, *z*) = *ϵδ*(*z*). Noting that 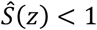, we have *J*(*t, z*) ≤ *w*(*t, z*) for any *t* > 0 and 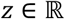 from the comparison theorem. The solution to (A48) is

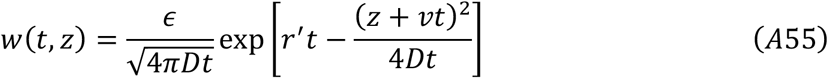

where *r*′ = *β*′ – (*γ* + *α*′). This follows by noting that *w*(*t, x*)*e*^−*r*′*t*^ follows a simple diffusion equation *∂w*/*∂t* = *D∂*^2^*w*/*∂x*^2^. By rearranging the exponents of (A49),

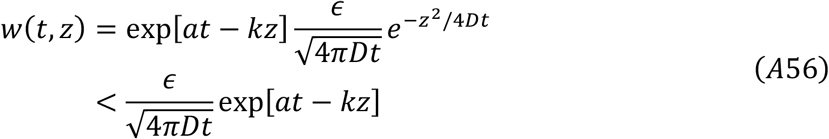

where

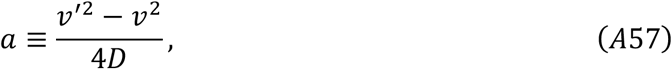

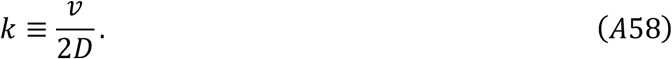

Here 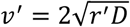 is the asymptotic wave speed if the mutant strain monopolizes the host population. Therefore, if *v*′ < *v*, then *a* < 0, and hence *w*(*t, z*) for a fixed *z* converges to zero as *t* goes to infinity; which in turn implies that *J*(*t, z*) converges to zero because *J*(*t, z*) ≤ *w*(*t, z*) for all *t* and *z*. Therefore, we conclude that any mutant that has a slower wave speed than the resident can never invade the population, implying that a strain that has the maximum wave speed 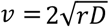 is locally evolutionarily stable.

